# Genomic and Transcriptomic Basis of Salinity Tolerance in Dry Pea

**DOI:** 10.64898/2026.05.05.722931

**Authors:** Shailesh Raj Acharya, Emmanuella Bredu, Harry Navasca, Hannah Worral, Lisa Piche, Rica Amor Saludares, Josephine Princy Johnson, Clarice Coyne, Kevin McPhee, Qi Zhang, Michael Ostlie, Nonoy Bandillo

## Abstract

Salinity is a major crop production constraint in dry pea (*Pisum sativum* L.), making the development of salt-tolerant varieties essential to improve crop productivity and land-use efficiency. The genetic mechanisms of salt tolerance in dry pea is largely unknown, and research on salt-tolerant genes is limited. In this study, we established comprehensive genomic and transcriptomic resources, along with a robust screening protocol, to dissect the genetic basis of salinity tolerance using two germplasm sets: the USDA pea diversity panel, consisting of approximately 200 globally sourced accessions, and a set of 300 modern elite lines from the NDSU Pulse Crops Breeding Program. Genetic variation for the salinity response was assessed based on ten phenotypic traits, with root dry weight, shoot dry weight, and specific root length identified as key indicators based on their heritability. Genome-wide association mapping uncovered significant genomic regions and several candidate genes linked to salt stress, with the strongest association found on chromosome 6. Overlapping QTL signals across traits suggest a shared genetic architecture underlying salinity tolerance. Field-based transcriptomic analysis further identified five putative genes involved in salinity response conserved across multiple crop species. Notably, Psat5g000800, encoding a glycosyl hydrolase gene, was markedly upregulated under salinity stress. These findings highlight the complex, multi-gene regulatory nature of salinity tolerance in dry pea and underscore the importance of functional validation of candidate genes. This study provides key insights and practical tools to support breeding efforts aimed at improving salt tolerance in dry pea.

## Introduction

Dry pea (*Pisum sativum* L.) is one of the most important legume crops, high in protein and carbohydrate content, and suitable for human and livestock consumption. It is the world’s second most widely grown pulse crop for protein. Dry pea performs well in neutral to slightly alkaline soils abundant in plant nutrients and beneficial microbes. However, changes in soil pH and elevated salinity levels can adversely affect productivity. Saline soils present major constraints to pea growth (Mustafa et al., 2019). Elevated levels of soluble salts, particularly chloride (Cl^-^) and sulfate (SO_4_^2-^) ions, generate excess osmotic stress that restricts water uptake by the plants (Alharbi et al., 2022). Salinity can also disrupt the ionic concentration within plant cells, resulting in cellular swelling and imbalances in ion homeostasis and osmotic regulation. Furthermore, saline stress can result in an overaccumulation of reactive oxygen species (ROS) which can affect molecular processes including protein and lipid degradation (dos Santos et al., 2022).

In the United States, North Dakota (ND) and Montana (MT) are the major dry pea-producing regions. In ND, at least 1.5 million acres are considered slightly saline, with salinity levels gradually increasing from moderate to high saline concentration over the years due to increasing annual precipitation and current agricultural practices (Scott, 2015). Dry pea has seen a significant reduction in overall seed yield to even moderate saline conditions compared to cereal and oilseed crops (Parihar et al., 2015; Page et al., 2021). The current field management practice to ameliorate the detrimental effects of salinity is to grow a salt-tolerant cover crop ahead of the dry pea crop and implement soil drying in the autumn to leach salts during spring snowmelt (Franzen et al., 2019). This highlights the importance of incorporating long-term genetic improvements to build on these cultural practices to help combat these effects and boost production in North Dakota as salinity levels continue to rise.

The understanding of salt-tolerance genetics in dry pea is complex and exacerbated by the dynamics of osmotic and ionic interaction during salinity stress (Parihar et al., 2015). Assessing the effect of salinity in dry pea at various developmental stages is crucial, as salt affects vegetative and reproductive stages differently due to variations in physiological and metabolic demands at each growth stage (Chaudhary et.al., 2024). This highlights the importance of evaluating dry pea plants at both vegetative and reproductive stages.

Salt toxicity can be caused by a mixture of salts that affect plant growth and development, and salt tolerance can vary at different molecular levels. For example, the salt overly sensitive (SOS) gene in *Arabidopsis* and the HKT1 gene in *O. sativa* encode sodium antiporters that remove excess sodium ions during salinity stress (Campbell et al., 2017).

Different osmolytes, like ROS in rice, wheat, and tobacco, also contribute to salinity resistance (Alkharabsheh et al., 2021; Li et al., 2024). MYB transcription factor signals salinity resistance in rice, while MAP kinases regulate cell shape during osmotic stress (Munns and Tester, 2008). MicroRNA expressions are enhanced in saline-tolerant maize and common beans (Chen et al., 2019). Studies in legume crops, mostly soybeans, have shown upregulation of antioxidant defense response genes (Jha et al., 2019; Hu et al., 2022). In pigeon pea, melatonin enhances stress tolerance by increasing flavonoids during salt stress (Song et al., 2022). This highlights the importance of studying salinity’s effect across multiple omics levels.

Dry pea production in ND is predominantly rainfed, where plants are exposed to complex and interacting abiotic and biotic stresses. Replicating these multifactorial field conditions in a greenhouse setting remains challenging. Consequently, most molecular studies to date have been conducted under controlled greenhouse environments, which do not fully capture interactions between salinity stress and other environmental factors, including extreme soil pH, temperature fluctuations, and metal toxicity. Nevertheless, controlled greenhouse studies provide a critical first step toward elucidating plant responses to salinity prior to evaluation under realistic field conditions.

In this study, we aimed to elucidate the genetic and molecular mechanisms underlying salinity tolerance in dry pea by identifying causal genetic variants and candidate genes associated with salt stress across contrasting environments. Initial evaluations were conducted under greenhouse conditions and subsequently compared with observations from field-grown lines. To achieve this, we integrated genomic and transcriptomic approaches: genome-wide analyses were used to map genetic variants associated with salinity stress and identify candidate loci, while RNA sequencing was employed to characterize differentially expressed genes in response to salinity. Integration of these datasets enabled the identification of key genes and pathways contributing to salinity tolerance. Collectively, these findings provide new insights into the genetic architecture of salinity tolerance in dry pea and identify valuable targets for breeding programs aimed at improving yield stability under saline conditions.

## Materials And Methods

### Plant materials

Two population sets were used in this study: (1) a USDA diversity panel, hereafter referred to as the USDA set, and (2) modern breeding lines from NDSU, hereafter referred to as the NDSU set. The USDA set comprised 193 accessions obtained from the USDA Western Regional Plant Introduction Program (Cheng et al., 2015; Bari et al., 2021), while the NDSU set included 300 modern elite lines from the North Dakota State University Pulse Crops Breeding Program (PCBP) (Bari et al., 2023). The commercial cultivar ‘Agassiz’ was included as a check. Altogether, approximately 500 accessions were evaluated, representing both public breeding lines and USDA germplasm accessions.

### Phenotypic screening under controlled greenhouse salinity treatment

This experiment was conducted in a controlled greenhouse environment in Fargo, ND. Seeds were initially germinated in a potting mixture for five days and then transplanted to cone-containers (556 mL; 6.4 cm diameter × 25.4 cm length) filled with a mix of perlite and sand (2:1, v:v). Containers were soaked in tubs filled with half-strength Hoagland solution (Li and Cheng, 2015) (∼ 2.5 cm above the bottom of the cone-containers) overnight and then transferred and maintained in tubs filled with tap water (∼ 2.5 cm above the bottom of the cone-containers) for 1 week (germination). Although germination rate was not measured as an independent trait, all seedlings that emerged were assessed for root and shoot related traits. After germination, the plants were split into two groups, one grown under non-stress conditions (i.e., tap water) and the other under salinity. Salinity stress was induced by setting the containers in the tubs filled with a Na_2_SO_4_ and MgSO_4_.7H_2_O mixture (2:1, M: M) at 10 dS m^-1^ about ∼2.5 cm deep. The tubs were filled with tap water (∼2.5 cm deep) for the non-stressed plants. Salt levels were progressively raised by 2.5 dS m^-1^ daily to prevent salinity shock, as previously described (Shahid et al., 2012). Once final salinity level was reached, plants remained under saline and non-saline conditions for four weeks, with solutions replenished weekly. Electrical conductivity (EC) and pH of the tub solutions were measured before and after replenishment using an EC/pH meter (Oakton PC 450, Oakton Instruments, Vernon Hills, IL) to monitor salinity levels.

The experiment used a split-plot design where the whole-plot treatment was the growing condition (saline vs. non-saline irrigation in the tubs) and the sub-plot treatment was the genotype. A total of four replications were used for the ∼200 accessions in the USDA set, and five replications were used for the 300 elite lines from the NDSU set under the aforementioned growing conditions, balancing statistical power and resource efficiency across populations. Data on Plant Height (Ht), Shoot Fresh Weight (SFW), Shoot Dry Weight (SDW), Root Dry Weight (RDW), Root to Shoot Dry Weight (RTSDW), Area Water Content (AWC), Total Above Ground Fresh Weight (TAGFW), Total Above Ground Dry Weight (TAGDW), Root Length (RL), and Specific Root Length (SRL) were sampled when the experiment was terminated (week 4). RDW and RTSDW measurements were recorded after oven-drying at 65°C for 48hrs. AWC and SRL were derived using the equation [AWC = TAAGFW – TAAGDW], SRL = [RL/RDW].

### Genotyping by sequencing

Leaf tissues from the greenhouse were harvested at different stages for the USDA set and the NDSU set. DNA from lyophilized tissues was extracted using DNAeasy Plant Mini Kit (Qiagen). Details on tissue collection and DNA extraction are provided in (Bari et al., 2021; Bari et al., 2023). Both the USDA set and the NDSU set were genotyped using genotyping-by-sequencing (GBS). Samples were sequenced using NovaSeq S1 × 100 (Illumina Technologies, USA). For the NDSU set, sequenced libraries with quality scores ≥ 30 were retained. For the USDA set, quality control was performed using FASTQC (Wingett and Andrews, 2018) and reads shorter than 50 bases were removed. High quality reads were aligned to the pea reference genome (Kreplak et al., 2019) and were analyzed using SAMtools (v1.10) (Li et al., 2009). SNP variants were called using FreeBayes (v1.3.2) (Garrison and Marth, 2012), which yielded a total of 380,527 SNP markers for the USDA set and 250,932 for the NDSU set. After removing SNPs > 90% missing values, heterozygosity >20%, and minor allele frequency < 5%, a total of 21,130 SNPs remained for the USDA set and 44,422 SNPs remained for the NDSU set. These high-quality SNPs were used for downstream analysis.

### Statistical analysis of phenotypic data

Best Linear Unbiased Predictions (BLUPs) were generated for all phenotypic values for both germplasm sets by fitting a mixed linear model (Equation 1) with the ASReml package in R (Butler et al., 2023)

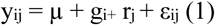

where y_ij_ is the observed phenotype, µ is the overall mean, g_i_ is the random effect of the i^th^ genotype, r_j_ is the effect of j^th^ replicate and ε_ij_ is the residual error which is assumed to be independent and identically distributed with a mean of zero and a common variance σ^2^ε.

Broad-sense heritability was calculated using Equation 2:

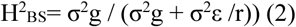

where σ^2^g represents genotypic variance, σ^2^ε represents residual or error variance and r represents number of replicates.

### Genome-Wide Association Mapping

Using the collected salinity data from the two germplasm sets and SNP datasets generated by genotyping-by-sequencing -- comprising 21,130 SNPs for the USDA set and 44,422 SNPs for the NDSU set -- we performed genome-wide association (GWA) mapping using a linear mixed model (Equation 3):

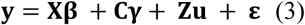

where y is a vector of the BLUPs, X, C and Z are design matrix for β, γ and u, respectively. β is a vector of fixed marker effects; γ is a vector of subpopulation effects; *u* is a vector of polygenic effects caused by relatedness
where u ∼ MVN (0, Kσ^2^u), ε is a vector of residuals where e ∼ MVN (0, Iσ^2^e). The matrix **K** is the realized relationship matrix estimated using the VanRaden Method (VanRaden, 2008). To control spurious association, population structure was accounted for by spectral decomposition of the genomic relationship matrix of the lines. The first three principal components were used in the standard GWA mixed linear model. The above model was implemented in GAPIT (Wang & Zhang, 2021). Association analyses were conducted within each population (e.g., NDSU set only, USDA set only). This is to avoid the confounding effects due to subpopulation structure that would arise by combining the two populations.

A comparison-wise error rate was calculated by the Li and Ji (2005) method to control the experiment-wise error rate. The effective number of independent tests (Meff) were calculated from the correlation matrix and eigenvalue decomposition of both SNP data within each population. The test criteria were adjusted with the following equation (Šidák,1967):

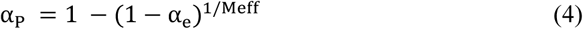

where, α_p_ is the computed comparison-wise error rate, and α_e_ is the trial-wise error rate. The experiment-wise error rate was controlled to limit Type I error arising from multiple testing by setting the adjusted significance level at αₑ = 0.05. Because SNPs are not fully independent due to long-range linkage disequilibrium in self-pollinated crops such as pea, the effective number of independent tests (Meff) was estimated and used to determine the genome-wide significance threshold.

Following GWA mapping, we defined a candidate region as 250kb region upstream and downstream of the most significant SNPs, based on the genome-wide estimated LD decay in the pea genome (Johnson et al., 2024). The region was defined to identify candidate genes located within these genomic intervals, with functional annotation obtained from *Pisum sativum* v1a genome assembly (Kreplak et al., 2019).

### Growth conditions and sample collection for RNA sequencing

RNA sequencing was conducted on four pea genotypes from the NDSU modern elite germplasm set, including the released cultivars ND Dawn yellow pea (Bandillo et al., 2021) and ND Victory (Bandillo et al., 2023), along with two advanced pre-release breeding lines. These genotypes were selected based on their contrasting phenotypic responses to salinity, making them well suited for gene expression analyses. The use of modern elite lines ensures direct relevance to commercial production, as growers prioritize salinity-tolerant cultivars adapted for field conditions.

Seed from these varieties was planted at the NDSU Carrington Research Extension Center in Carrington, ND, at a 40 x 1000 ft site selected for its natural saline gradient. This field was divided into 5 sections representing differing salinity levels (Treatment 1 = lowest, Treatment 5 = highest), which served as main plots within a split-plot design wherein each genotype was replicated twice per salinity treatment, serving as subplots and arranged left to right within each salinity level.

Leaves and root tissues at vegetative (V4: fourth true leaf has unfurled at the third node) and reproductive (R2: first open flower at one or more node) stages (Grains Research and Development Corporation, 2018), were collected from the field, stored in dry ice, and immediately frozen at -80°C for RNA extraction. Salinity measurements were conducted using a soil probe for both leaf and root tissues at the V4 and R2 stages. V4 and R2 stages were selected for evaluation because salinity stress is known to exert significant effects at these two developmental time points in dry pea (Chaudhary et.al., 2024).

### RNA extraction and sequencing

Total RNA was extracted from 200 mg of flash frozen tissue from root and leaf samples using theZymo^TM^ Quick RNA Plant Miniprep Kit (Zymo, USA) and following the manufacturer’s protocol. RNA quantification and quality was evaluated using a Qubit v4 Fluorometer (ThermoFisher Scientific, Waltham MA) and was also visualized using agarose gel electrophoresis. The RNA Integrity Number (RIN) value was checked using an Agilent 2100 Bioanalyzer System (Agilent Technologies, CA, USA) to ensure all samples passed the RIN threshold (RIN ≥ 8.0). The 3’ RNAseq libraries were prepared from ∼500ng of total RNA per sample using the Lexogen QuantSeq 3’ mRNA-Seq Library Prep Kit FWD for Illumina (Corley et al., 2019). The libraries were quantified on a Molecular Devices Spectra Max M2 plate reader (with the intercalating dye QuantiFluor) and pooled accordingly for maximum evenness. The pool was quantified by digital PCR and sequenced on an Illumina NextSeq500 sequencer and de-multiplexed based upon six base i7 indices using Illumina bcl2fastq software (version 2.20) (Illumina, 2024).

### RNA-Seq quantification and mapping

RNA samples with fewer than 200,000 de-multiplexed reads were excluded from further analysis. For the remaining samples, Illumina adapters on the 3’ end and the first twelve bases on the 5’ end (corresponding to the random primer) were removed from the de-multiplexed fastq files using Trimmomatic (version 0.36) (Bolger et al., 2014). Poly-A tails and poly-G stretches of at least 10 bases in length were then removed using the BBDuk program from the package BBMap (version 37.50) (Bushnell, 2014), keeping reads at least 18 bases in length after trimming. Poly-G stretches result from sequencing past the ends of short fragments (G = no signal). The trimmed reads were aligned to version 1a of the *Pisum sativum* genome assembly (Kreplak et al., 2019) using the STAR aligner (version 2.7.10b) (Dobin et al., 2013). For the STAR indexing step, the gff3 annotation file was converted to gtf format with the gffread program (version 0.10.4) from cufflinks (Trapnell et al., 2013). The output SAM files were converted to BAM using SAMtools (version 1.15.1), and the number of reads overlapping each gene in the gff3 file on the forward strand were counted using HTSeq-count (version 0.6.1) (Anders et al., 2015).

The R package DESeq2 (version 1.36.0) (Love et al., 2014) was used to obtain both normalized and variance-stabilized counts, to conduct principal components analyses of the 500 most variably expressed genes after count normalization and variance stabilizing transformation, and to produce heat maps of distances between clustered samples. For the variance-stabilized counts, genes with fewer than two counts per sample on average were excluded. Counts were produced for all samples and for the leaf and root samples separately. For the leaf and root samples, additional gene count matrices were produced after excluding samples with fewer than 10K (leaf) or fewer than 6K (root) genes expressed with a minimum count of 10. These cutoffs were chosen based upon visual inspection of the relevant histograms. Tests for differential expression (DE) were performed for the leaf and root samples separately (i.e., independent DESeq2 analyses were performed for leaf and root). Only genes that survived the filter for low expression (in each tissue) were included in the DE tests. For the remaining leaf samples after sample filtering, all pairwise contrasts between the five treatments (10 contrasts) were performed within each developmental stage (i.e., within LeafR2 and LeafV4) (10 contrasts within each stage = 20 contrasts). In addition, a test for DE between the two stages, LeafR2 and LeafV4, was performed. The same approach was then applied to the root samples. All DE tests were performed with and without Bayesian shrinkage of the log fold change (LFC) estimates via the “apeglm” method (Zhu et al., 2019). DE for sample can be summarized as: 2 tissues x (2 stages x 10 contrasts + 1 between stage contrast) x 2 statistical methods = 84 DE tests in total.

Gene Enrichment Analysis for genes upregulated from DeSeq was performed using ShinyGo v0.8 (Ge et al., 2020) and tested against the pea reference transcriptome. All significant GO terms related to upregulated genes were reported and upregulated genes were categorized based on their functional involvement. Fold enrichment was calculated as the total percentage of upregulated genes belonging to a pathway divided by total background gene supplied. Higher fold values show that a particular GO term is over-represented in the upregulated DEGs. GO terms were categorized into biological processes, molecular functions and cellular components, with enrichment significance assessed using the hypergeometric test method and false discovery rate (FDR) correction at an adjusted p-value threshold of 0.05.

## Results

### Salinity stress reduces biomass and alters root elongation traits

Phenotypic trait performances were evaluated under salinity stress and non-saline conditions. Both the USDA set and the NDSU set showed significant changes across all ten measured traits under salinity stress. The measured phenotypic traits decreased significantly in the saline environment, except for Root to Shoot Dry Weight (RTDSW) and Specific Root Length (SRL). Plant height decreased by 15%, suggesting a decline in vertical growth. Biomass-related traits also decreased by around 30% for Shoot Fresh Weight (SFW), Shoot Dry Weight (SDW) and Root Dry Weight (RDW) (Fig1, Table S1). Conversely, SRL increased by 20% on average in both populations, resulting from a higher reduction in RDW than RL under salt stress.

**Fig 1:**
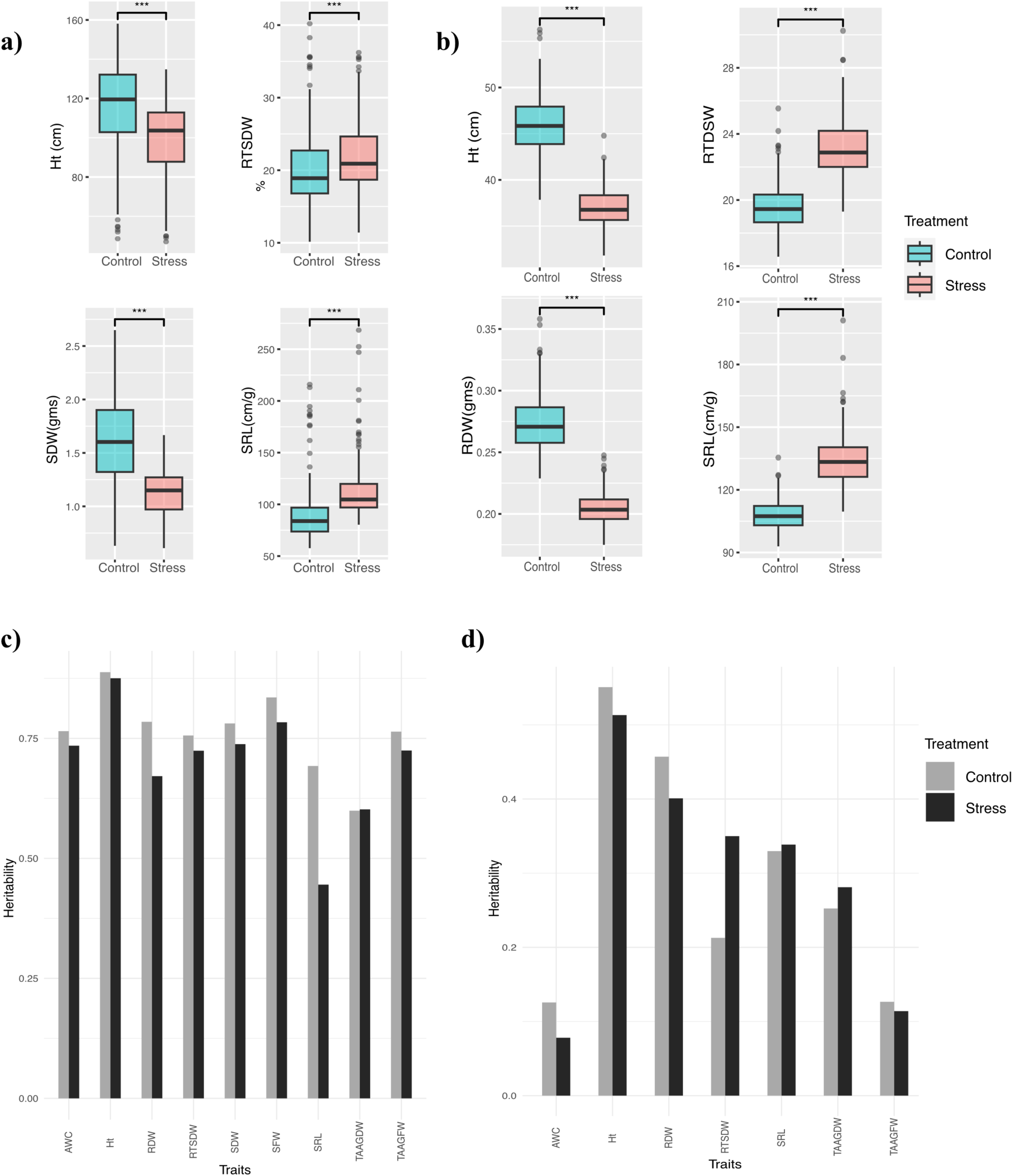
Phenotypic variations measured across traits. Phenotypic variations measured across a) USDA set b) NDSU set. Bar plot shows traits measure under non-stressed (control) and salt stressed conditions. Asterisks on top of bar plot shows significant mean differences among control and stressed phenotypes. Broad-sense Heritability measured in c) USDA set d) NDSU set. The height of the bar plot shows heritability estimates for different traits under control and salt-stress conditions. AWC: Area Water Content, Ht: height, RDW: Root Dry Weight, SDW: Shoot Dry Weight, SFW: Shoot Fresh Weight, SRL: Specific Root Length, TAAGDW: Total Area Above Ground Dry Weight, TAGGFW: Total Area Above Ground Fresh Weight, RTDSW: Root to Shoot Dry Weight

### Biomass related traits exhibited higher genetic variation in the USDA set compared to NDSU set

Broad-sense heritability for the USDA set ranged from 0.44 for SRL to 0.88 for height, while heritability for the NDSU set ranged from 0.07 for AWC to 0.55 for height. Heritability values were higher overall for the USDA set than the NDSU set, and apart from RTSDW, SRL, and TAAGDW in the NDSU set, were higher under control conditions than under stress conditions (Fig 1).

### Genome-wide association mapping identifies potential salinity stress regulation genes

Genome-wide association mapping was conducted separately for the USDA and NDSU germplasm sets. Analyses under salinity stress conditions were used to identify SNPs associated with stress-responsive trait variation, whereas analyses under control conditions captured baseline genetic variation independent of stress.

GWA mapping of the USDA panel, using 21,130 SNPs across nine phenotypes, identified 15 highly significant SNPs associated with height, RTSDW, SDW, and SRL. SRL showed the strongest association (P ≈ 10⁻⁹), explaining 19% of the phenotypic variance (PVE), while other traits exhibited PVEs ranging from 6% to 20% (Table S2). In the NDSU panel, GWA mapping with 44,422 SNPs across the same nine phenotypes identified 30 significant SNPs. The strongest association was observed for RDW (19% PVE), whereas associations for height, RTDSW, and SRL peaked at ∼10% PVE, consistent with the involvement of multiple small-effect loci and/or environmental influences under saline conditions.

To assess whether the most significant GWA SNPs localized to functionally relevant regions, genes annotated within a 250-kb window flanking each SNP were examined (Fig. 2). This analysis identified 39 genes in the USDA set and four genes in the NDSU set. In the USDA panel, most candidate genes were associated with SDW. Notably, a 4.4-kb gene annotated as a major sigma-70 factor (Sigma factor B) spans SNP S5LG3_1667684 on chromosome 5 (Table 1). Sigma factors regulate RNA polymerase activity and play key roles in abiotic stress responses by controlling stress-responsive gene expression. In addition, a 3.7-kb gene, Psat6g207360, encoding a Frigida-like protein (FLP), spans two SNPs on chromosome 6 and is associated with plant height. FLPs are implicated in abscisic acid (ABA) signaling pathways that mediate ion homeostasis under salinity stress.

**Fig 2:**
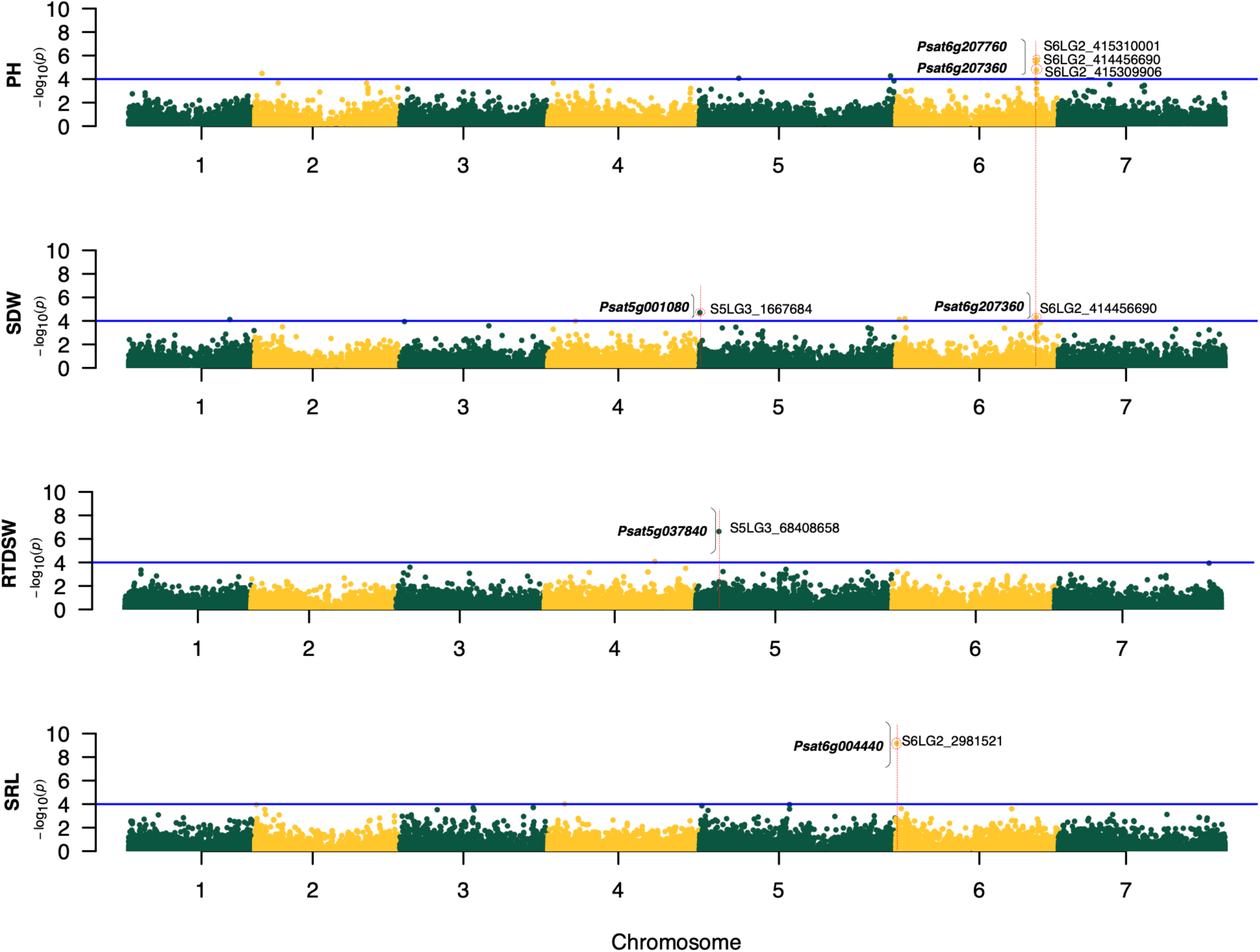
Manhattan plots of height and biomass related traits for USDA set. Threshold value represented by solid blue horizontal line is obtained by Li and Ji method ≈ 4. Dotted red circle represents most significant GWA mapping SNP crossing this threshold. Candidate genes spanning the associated SNP are represented in bold letters, with brackets separating the encircled SNP. PH: Plant Height, SDW: Shoot Dry Weight, RTDSW: Root to Shoot Dry Weight, SRL: Specific Root Length

**Table 1:**
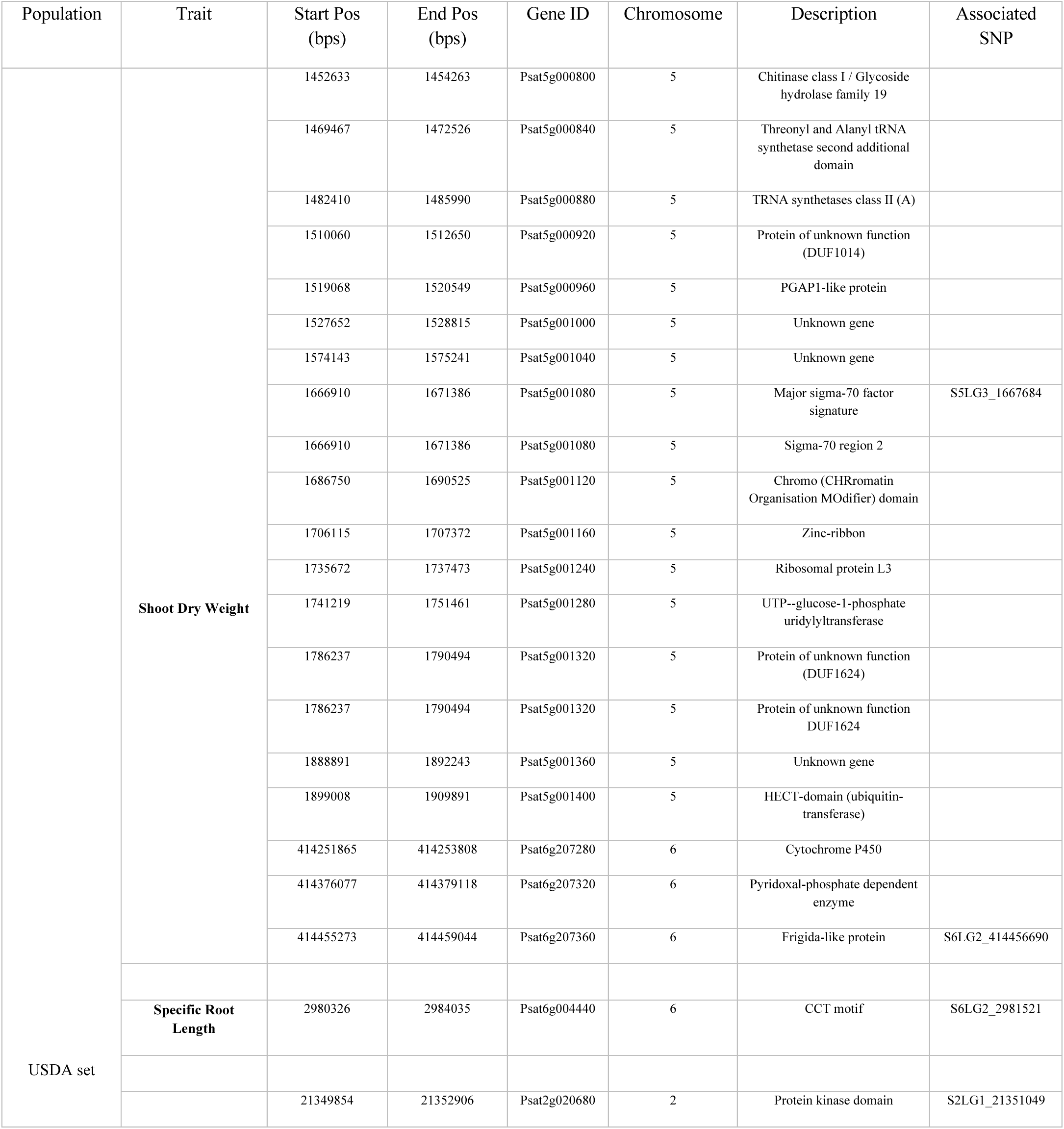
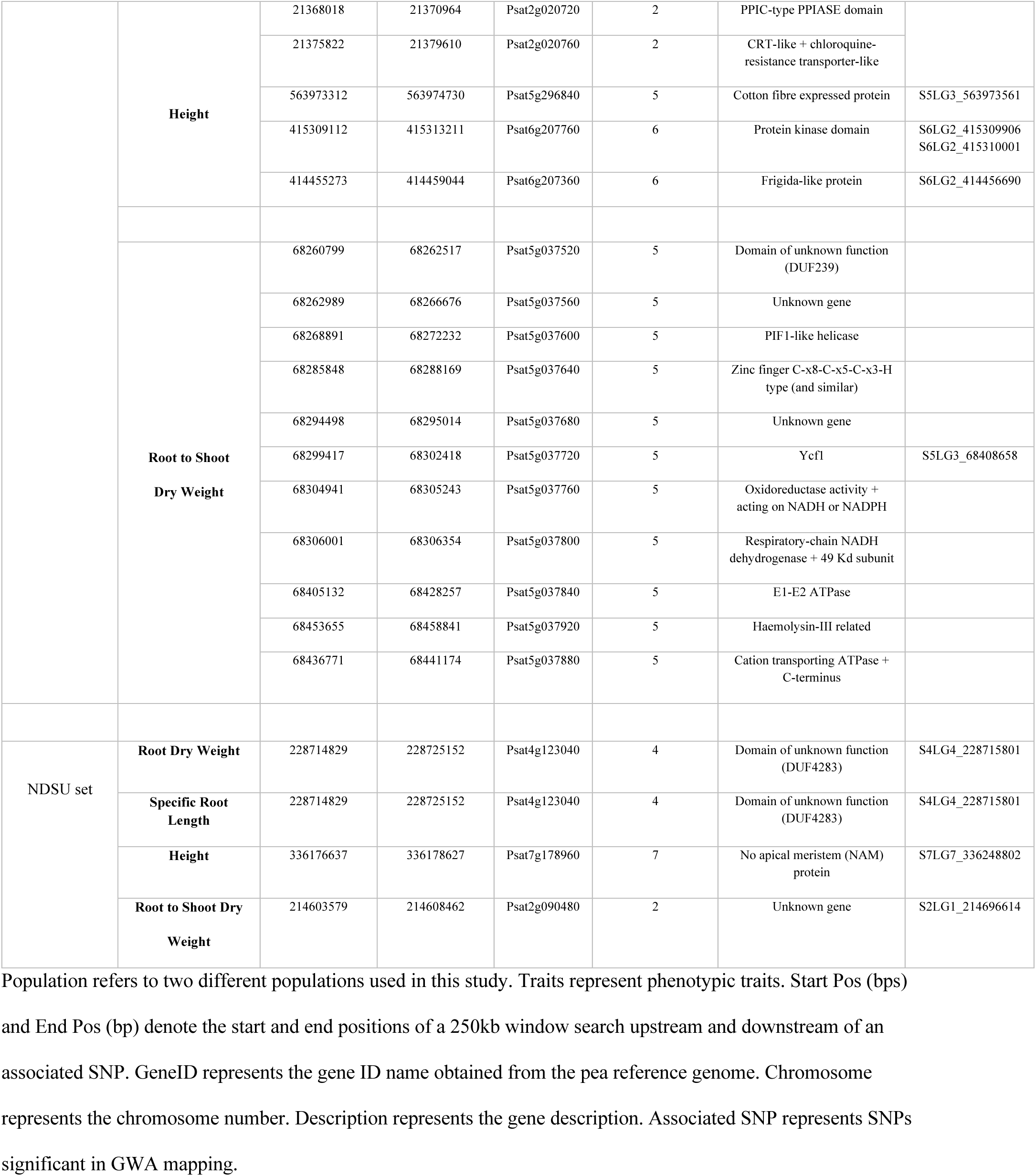
Summary of GWA mapping identified SNPs, associated genes and their functions.

A height-associated GWA SNP also overlapped a 2.9-kb gene, Psat2g020720, annotated as a PPIC-type peptidyl–prolyl isomerase (PPIase), spanning SNP S2LG1_21351049. PPIases primarily regulate protein folding and stability under stress conditions. For RDW, a 23-kb gene, Psat5g037840, annotated as an E1–E2 ATPase, spanned SNP S5LG3_68408658 on chromosome 5; this ATPase is involved in sodium and potassium transport, contributing to ionic homeostasis during salt stress.

In the NDSU panel, GWA mapping identified four candidate genes, three of which have unknown functions and span SNP S4LG4_228715801 (Fig. S1). The remaining candidate was a 1.9-kb gene, Psat7g178960, annotated as a No Apical Meristem (NAM) protein, spanning SNP S7LG7_336248802 on chromosome 7 and associated with height. NAM proteins regulate developmental processes and mediate cell apoptosis in response to salinity stress.

### A field-based transcriptome analysis identified highly upregulated genes under salinity stress

To further investigate the genetic mechanisms underlying salinity stress responses and to complement the GWA mapping results from the two sets, we conducted a field-based transcriptomic analysis using varieties and advanced lines with contrasting salinity tolerance. Plants were grown under five salinity treatments in the field (Fig. 3A), spanning a conductivity range from 114 to 4,910 µS/cm.

**Fig 3:**
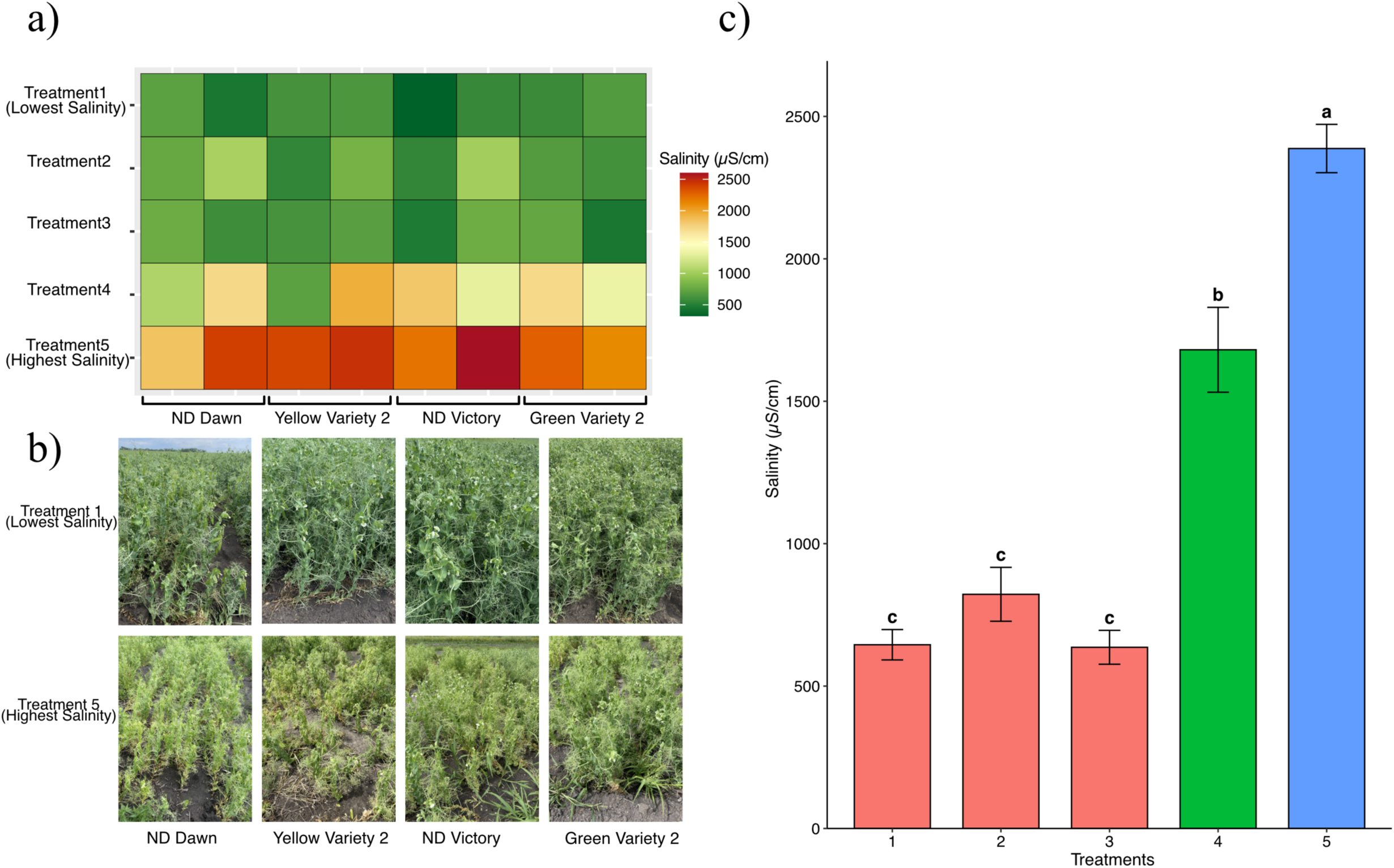
Salinity levels across experimental field plots. a) Heatmap of experimental plots shows differences in soil salinity level. b) Experimental plots show the impact of high salinity on pea growth and development, with salinity increasing progressively from Treatment 1 (lowest) to Treatment 5 (highest). c) The average salinity levels were significantly higher in treatments 4 and 5 than in treatments 1, 2, and 3. Treatment 5 had the highest salinity among all treatments while treatments 1,2 and 3 did not show any statistical significance to salinity levels.

Based on field salinity measurements (Treatment 1 = lowest salinity; Treatment 5 = highest salinity), the original five treatments were consolidated into three levels: Treatments 1, 4, and 5. This consolidation was justified because Treatments 1, 2, and 3 did not differ significantly and were therefore combined into a single category (Treatment 1) (Fig. 3C). Pooling these treatments increased the effective sample size and, consequently, the statistical power of the analysis. The average salinity of the highest treatment (Treatment 5) was 2,387.3 µS/cm, whereas the lowest treatment (Treatment 1) averaged 411.91 µS/cm (Fig. 3A). Treatment 1 served as the baseline for RNA sequencing of root and leaf tissues collected at the vegetative (V4) and reproductive (R2) stages.

Differential gene expression (DEG) analysis from RNAseq data revealed higher overall gene expression in treatment 5 across both roots and leaves, except in R2 leaf tissue, where treatment 4 showed a higher number of DEGs (Fig S2). Upregulated DEGs (log_2_ fold change ≥ 2) totaled 625 in leaves and 771 in roots. The highest count (427) was in V4 root tissue under treatment 5, while the lowest (62) was in R2 leaf tissue under treatment 4 (Fig 4). At V4, 86 and 43 upregulated genes were shared between treatments 4 and 5 in leaves and roots, respectively, with 42 and 32 common at R2. Five genes were common across all stages and treatments (Fig S4). Two of these, Psat0s1036g0040: Natural resistance-associated macrophage protein and Psat0s4771g0040 Prolyl oligopeptidase + N-terminal beta-propeller domain, are directly involved in salinity stress response (Table 2).

**Fig 4:**
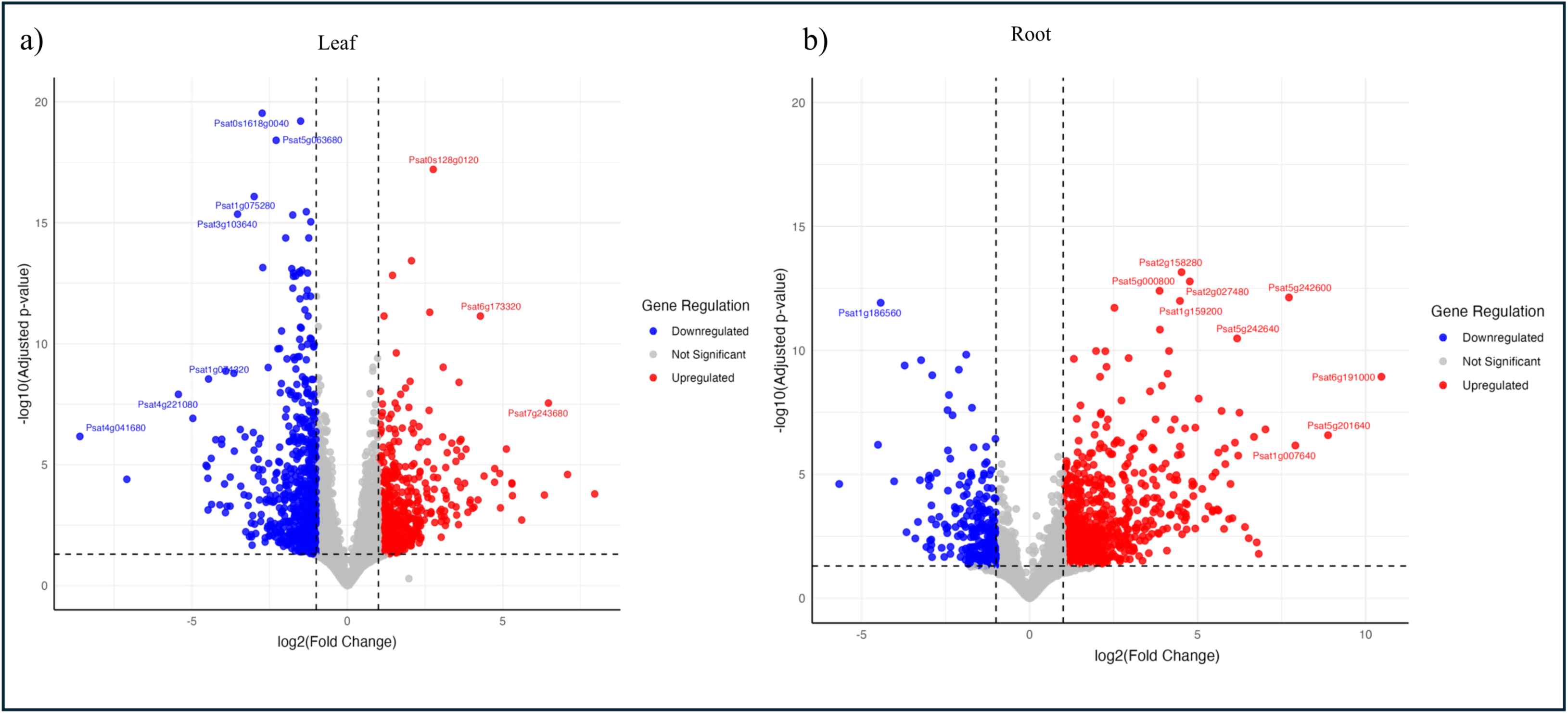
Volcano plot of differentially expressed genes across tissues. Volcano plot of differentially expressed genes at the highest salinity treatment (Treatment 5) for a) Leaves and b) Roots. Genes with a log_2_foldchange ≥ 2 threshold were considered upregulated.

**Table 2:**
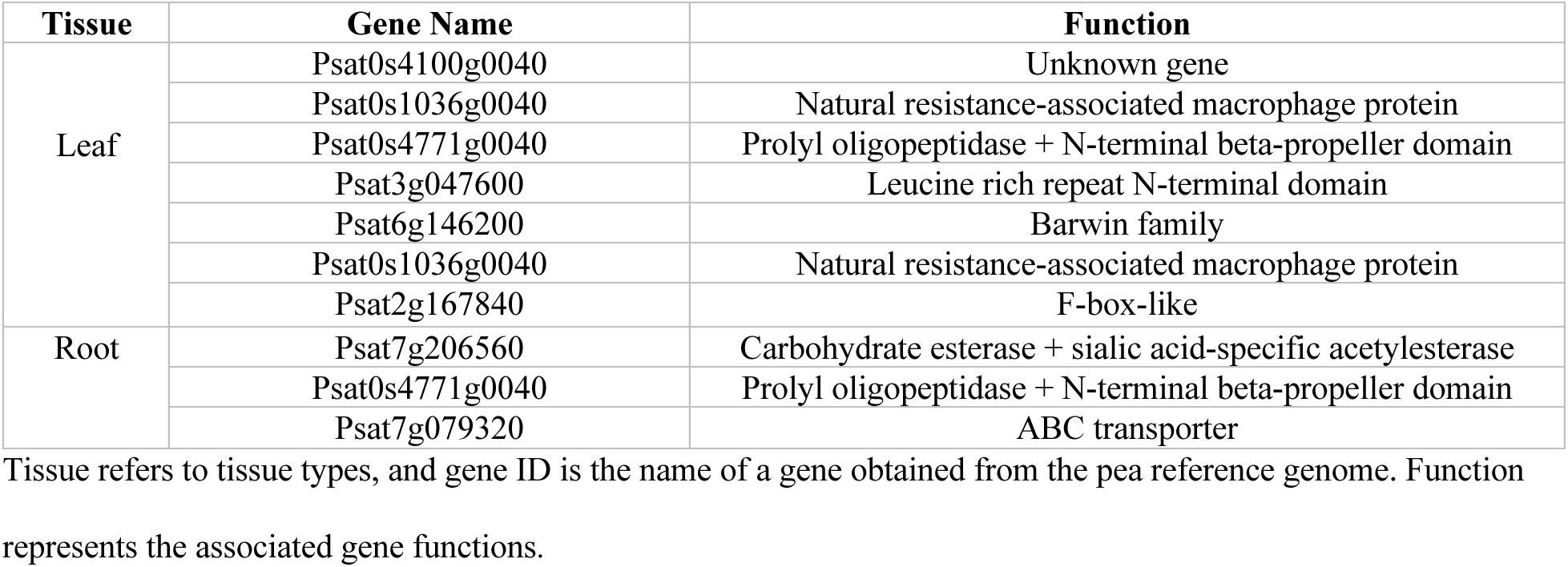
Common upregulated DEGs across leaf and roots.

Gene Ontology (GO) enrichment analysis of upregulated DEGs identified key biological processes, pathways, and molecular functions involved in stress regulation. In leaf tissue at V4-treatment 5, the highest number of upregulated DEGs were linked to plant hormone signal transduction (Pathway ID - Psat04075) (Fig S5), including ‘Psat2g169400’ and ‘Psat4g175160’, associated with the GH3 auxin-responsive promoter. Four DEGs were enriched in alpha-linolenic acid metabolism (Pathway ID - Psat00592), linked to reactive oxygen species production. Additionally, six upregulated heat shock proteins (Hsps) were enriched in protein processing in the endoplasmic reticulum (Pathway ID - Psat04141) (Table S3 and S4). The highest fold enrichment was observed in Betalain biosynthesis (Pathway ID - Psat00965), with four DEGs involved in betaxanthin production. Glucosinolate biosynthesis (Pathway ID - Psat2g114240) showed the second highest enrichment, with five DEGs linked to salinity stress.

At the R2-treatment 4 stage, photosynthesis (Pathway ID - Psat00195) was enriched with six DEGs, and arginine and proline metabolism (Pathway ID - Psat00330) involved four DEGs. In root tissue at V4-treatment 5, glucosinolate biosynthesis was enriched with two DEGs. The highest number of upregulated DEGs in root tissue at V4-treatment 4 and V4-treatment 5 were linked to the biosynthesis of secondary metabolites (Pathway ID - Psat01110). Pentose and glucuronate interconversions (Pathway ID - Psat00040) were enriched in V4-treatment 4 with two DEGs. At root R2-treatment 5, phenylpropanoid biosynthesis (Pathway ID - Psat00940) and the MAPK signaling pathway (Pathway ID - Psat04016) were highly enriched, each with five DEGs (Table S4). No GO-enriched terms were observed for root R2-treatment 4. A complete list of all GO enrichment term and their associated genes are presented in Table S4.

### A potential stress regulation gene identified from overlapping GWA mapping and RNA-seq studies

A common gene, Psat5g000800, was present among the differentially upregulated genes in RNAseq with GWA mapping functional analysis, which was associated with shoot dry weight. Psat5g000800 is a 1.6kb long chitinase-related gene belonging to the glycosyl hydrolase (GH) family. The GH family is comprised of diverse enzymes that hydrolyze glycosidic bonds, playing key roles in cell wall remodeling, osmotic adjustment, and signal transduction under salinity stress primarily for cellular homeostasis, ionic balance and reactive oxygen species (ROS) scavenging. Interestingly, our GO analysis revealed that critical pathways involved in glycosyl hydrolase-mediated stress adaptation, such as phenylpropanoid and MAPK signaling pathways, were significantly enriched. This highlights the importance of GH genes in stress responsiveness to salinity.

## Discussion

This study integrated genomic and transcriptomic approaches to identify key candidate genes associated with salinity tolerance in dry pea. Our genomic analysis, based on GWA mapping mapped genetic variants linked to salinity stress responses. Complementing this, RNA sequencing enabled the identification of differentially expressed genes enriched under salinity stress across multiple tissues and developmental stages. Through this integrated approach, several functionally relevant candidate genes were identified, providing potential targets for improving salinity tolerance in dry pea.

GWA mapping results highlighted key genes involved in salinity stress regulation. Notably, a zinc finger transcription factor, CONSTANS-Like (CCT) gene annotated as *Psat6g004440*, was associated with specific root length (SRL). This gene likely plays a role in environmental stress response and is homologous to *GmCOLa* in transgenic soybean, which confers enhanced salt stress tolerance (Xu et al., 2023). In chickpeas, Garg et al. (2016) reported that CCT genes were differentially expressed under salinity stress, indicating a potential role in coordinating stress signaling with developmental pathways. Additionally, CCT family genes have also been reported in regulating abiotic stress responses across various crops, including cotton (Qin et al., 2018), sunflower (Niu et al., 2022), *Petunia* (Khatun et al., 2021), *Arabidopsis* (Min et al., 2015), and *Andrographis paniculata* (Zhao et al., 2025), highlighting their conserved functional role in stress adaptation. Similarly, *Psat5g001080* gene, encoding a major sigma-70 factor/RNA polymerase regulatory protein, was associated with shoot dry weight (SDW). Sigma factors modulate RNA polymerase activity and activate stress-responsive gene networks during abiotic stress. Chaudhari et al. (2020) noted a significantly higher abundance of genes encoding the sigma-70 factor in peas under highly acidic conditions, which helps shape their rhizosphere microbiome for stress tolerance. Likewise, sigma-70-mediated molecular responses to drought and salinity stress have also been reported in chickpea (Garg et al., 2016). A functionally similar gene in *Arabidopsis*, *SOLDAT8*, is known to regulate singlet oxygen-induced programmed cell death, which restricts oxidative stress propagation (Coll et al., 2009).

Although salinity tolerance has been widely studied in various crops (Genc et al., 2007; Møller et al., 2009; Jha et al., 2019; Giordano et al., 2021; Hu et al., 2022; Afzal et al., 2023), its underlying molecular mechanisms in dry peas remain poorly understood. Our transcriptomic analysis revealed that salt-tolerant genes exhibited elevated expression across multiple growth stages and tissues. DEGs encoding natural resistance-associated macrophage proteins (NRAMPs) were consistently expressed in both root and leaf tissues under all treatments. NRAMP transporters are critical for maintaining ion homeostasis under salinity stress, in addition to their established roles in heavy metal detoxification (Ishimaru et al., 2012; Yan et al., 2024; Yu et al., 2025; Toyokura et al., 2025). In soybean, several NRAMP genes were differentially regulated under nutrient stress, which share signaling crosstalk with different abiotic stresses including salinity stress (Qin et al., 2017). Similarly, prolyl-oligopeptidase proteins differentially expressed in our study correspond to those reported in *Arabidopsis,* where they enhance proline accumulation and confer salt tolerance (Sun et al., 2013). Additionally, increased levels of prolyl-oligopeptidase-like proteins and enzymes involved in proline metabolism have been reported in salt-stressed chickpea, suggesting their involvement in salinity tolerance pathways (Arefian et al., 2019).

Gene Ontology (GO) enrichment analysis of upregulated genes further confirmed the involvement of critical stress-response pathways. These included genes encoding phosphatase-2C and UDP-glycosyltransferases involved in glucosinolate biosynthesis, as well as key components of the MAPK signaling cascade linked to abscisic acid (ABA) regulation and MYB transcription factor-mediated ion homeostasis (Parihar et al., 2015; Jha et al., 2019). In chickpeas, Singh et al. (2018) reported several MAPK genes that are rapidly induced by abiotic stresses and ABA signaling, while MYB transcription factors linked to ion homeostasis under salt stress have also been reported (Duan et al., 2025; Kaashyap et al., 2018, 2022), confirming their role in stress adaptation. Additionally, comparative transcriptomic analyses have revealed that salt tolerance is heavily dependent on the integration of ABA signaling and MAPK cascades to regulate downstream stress-responsive genes (Sinha et al., 2011).

In our study, photosystem I (PSI) and photosystem II (PSII) genes were upregulated under salinity stress, a pattern that aligns with earlier findings in chickpeas, where salt-tolerant varieties showed higher expression of PSI and PSII genes, contributing to chlorophyll stability and reducing structural damage to chloroplast under salinity stress. (Khan et al., 2023). Additionally, similar upregulation of PSI and PSII genes has been reported in leaf tissues of other plant species, indicating an early photosynthetic response of the photosystem to salt stress (Zhang et al., 2020). Our study also showed several upregulated heat shock proteins (Hsp20/70/90) under salinity stress, indicating activation of molecular chaperon systems involved in preventing protein aggregation. Comparable results have been reported in salt-stressed chickpea, where the Hsp70 and Hsp90 families were strongly induced under salinity stress (Khan et al., 2023; Arefian et al., 2019), while in common bean, Hsps’ contributed to broad abiotic stress resistance (Büyük et al., 2016). Furthermore, our study demonstrated that cytochrome P450 enzymes were upregulated under salinity, consistent with chickpea studies showing their involvement in oxidoreductase activity, stress adaptation pathways, membrane stability and iron binding capacity (Kumar et al., 2021; Garg et al., 2016). Additionally, the upregulation of several peroxidases, including glutathione peroxidase, further highlights the importance of oxidative stress mitigation under salinity stress, consistent with findings in chickpea (Arefian et al., 2019), pea (Farooq et al., 2021) and soybean (Jin et al., 2019).

Among the genes identified through both GWA mapping and transcriptomic analysis, *Psat5g000800* gene, encoding an endochitinase A2 belonging to the glycosyl hydrolase 19 (GH19) family, emerged as a strong candidate associated with salinity tolerance. Members of the GH19 family are widely linked with salicylic acid (SA)-mediated plant defense responses (Upadhyay, 2024). *Psat5g000800* gene identified in the present study shares homology to chitinase-related proteins, which are known to be SA-responsive in legumes (Egorova, 2024). Beyond their classical role in pathogen defense, chitinases have been increasingly recognized for their involvement in abiotic stress responses, where they contribute to osmotic adjustment and protection against oxidative damage (Kaashyap et al., 2018). In pea, chitinase-related gene have been shown to function as osmoprotectants, mitigating reactive oxygen species (ROS)-induced cellular damage under salinity stress (Ahmad et al., 2022). Transcriptomic studies in chickpea have also reported the induction of defense related glycosyl hydrolase family genes under stress, including chitinase-like proteins associated with ROS detoxification and stress adaptation (Irum et al., 2025). Although specific GH19 endochitinase A2 homologs induced under salinity stress have not been extensively characterized, available evidence suggests a conserved role for chitinase-related protein in mediating oxidative stress responses and cellular protection under saline conditions. Collectively, these findings suggest that *Psat5g000800* gene may play a dual role in coordinating defense signaling and osmotic stress mitigation and its significant association with shoot dry weight (SDW) trait further highlights its potential as a candidate gene for improve salinity tolerance in pea breeding.

A secondary objective of the study was to identify high-priority, heritable traits suitable for targeted breeding. Trait selection was based on data from both the USDA set and the NDSU set. The selection criteria incorporated agronomic relevance, genetic signal strength from GWA mapping, and functional biological evidence. Four traits were prioritized, of which RDW, SDW, and SRL demonstrated consistently high heritability across experiments, making these traits as robust targets for breeding. Furthermore, the study was structured to closely simulate the real-world application of identified genes within an applied public breeding program. The breeding materials used were unstructured elite lines derived from multiple biparental crosses involving modern elite parents. This population captured the genetic diversity typical of an advanced breeding pipeline, and the selected lines were both available and relevant to ongoing selection efforts. This alignment ensured that the study’s results could be directly integrated into selection decisions and demonstrated the feasibility of applying gene discovery and expression profiling to improve stress resilience in pulse crops.

## Conclusion

In conclusion, this study elucidates the genomic and transcriptomic foundations of salinity tolerance in dry pea and identifies elite breeding lines and germplasm with promising salt tolerance for potential future release. By integrating genome-wide variation with gene expression analyses, we provide new insights into the molecular mechanisms governing salt stress responses. Functional validation of the candidate genes identified here will be essential to confirm their roles in stress adaptation and to accelerate the development of salt-tolerant dry pea cultivars through targeted breeding strategies.

## Statements and Declaration

## Acknowledgments and Funding

The authors would like to acknowledge the funding provided by the North Dakota Department of Agriculture through the Specialty Crop Block Grant Program (Award no. 18-265 and 19-429), and the Northern Pulse Growers Association for continued funding support on development of the NDSU advanced pea breeding lines. The authors would also like to acknowledge the funding provided by USDA Plant Genetic Resource Evaluation Grant for the GS analysis, USA Dry Pea and Lentil Council Research Committee for the field phenotyping, and USDA ARS Pulse Crop Health Initiative for the SNP genotyping and support from USDA ARS Project: 5348-21000-017-00. This investigation used resources from the Center for Computationally Assisted Science and Technology (CCAST) at North Dakota State University, Fargo, ND, USA, which were made possible in part by NSF MRI Award No. 2019077.

## Author Contributions

**SA**: Data curation; formal analysis; investigation; methodology; writing–original draft; writing–review and editing. **EB**: Investigation, writing review and editing. **HN:** Investigation, writing review and editing. **HW**: Writing review and editing. **LP:** Investigation, writing review and editing. **RAS:** Investigation; methodology. **JPJ**: Investigation; methodology. **CC:** Writing review and editing. **KM:** Writing review and editing. **QZ:** Investigation; methodology; writing–review and editing. **MO:** Conceptualization; investigation; methodology; writing–review and editing. **NB:** Conceptualization; funding acquisition; investigation; methodology; project administration; supervision; writing–review and editing.

## Data availability

The SNP dataset utilized in this study is accessible through https://www.ncbi.nlm.nih.gov/sra/PRJNA730349. For access to the phenotype data, please contact the corresponding author.

