## Supplemental Figures for "Genomic and Transcriptomic Basis of Salinity Tolerance in Dry Pea"

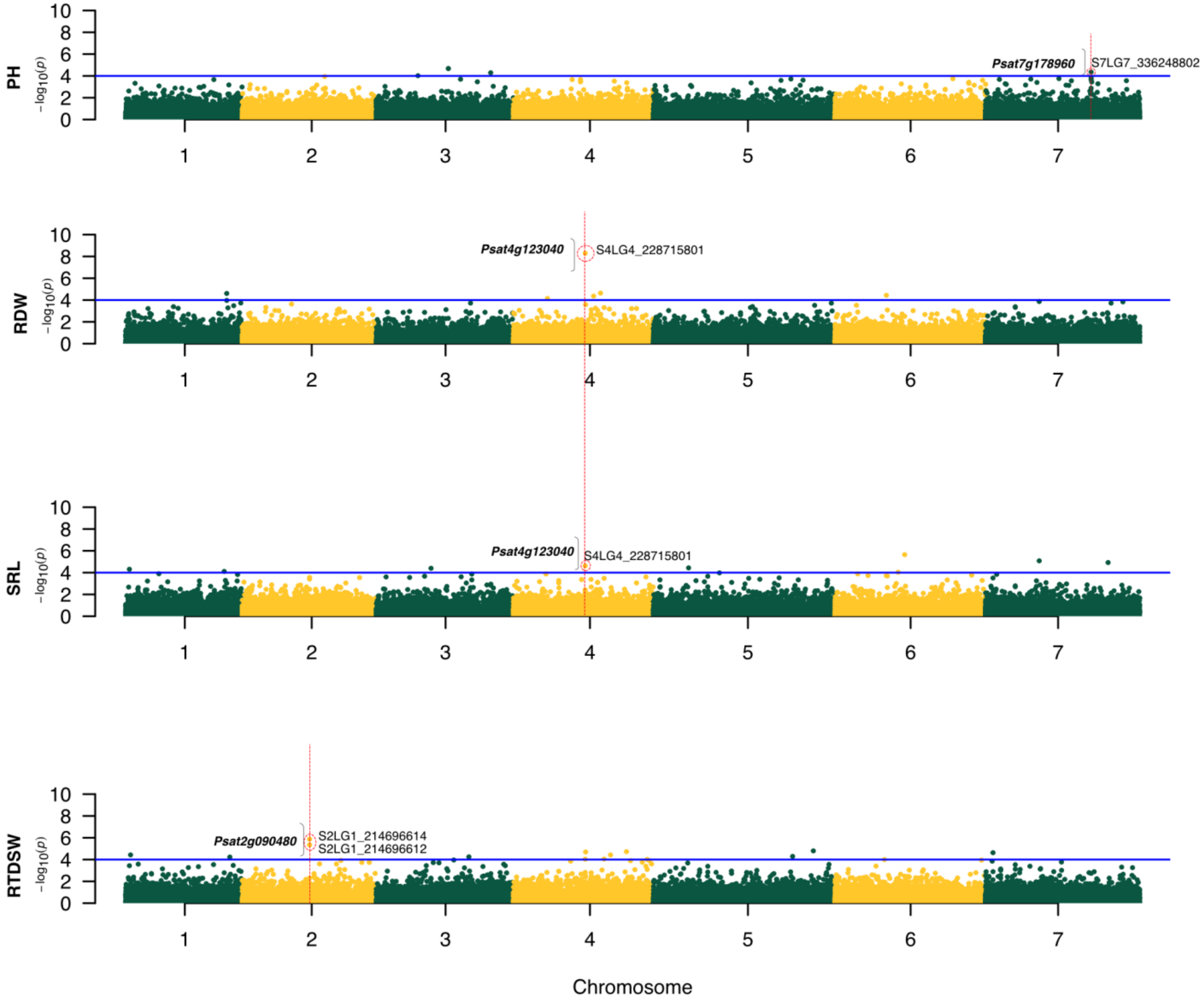


**Fig S1:** Manhattan plots of height and biomass related traits for NDSU germplasm panel. Threshold value represented by solid blue horizontal line is obtained by Li and Ji method ≈ 4. Dotted red circle represents most significant GWAS SNP crossing this threshold. Candidate genes spanning the associated SNP are represented in bold letters, with brackets separating the encircled SNP.

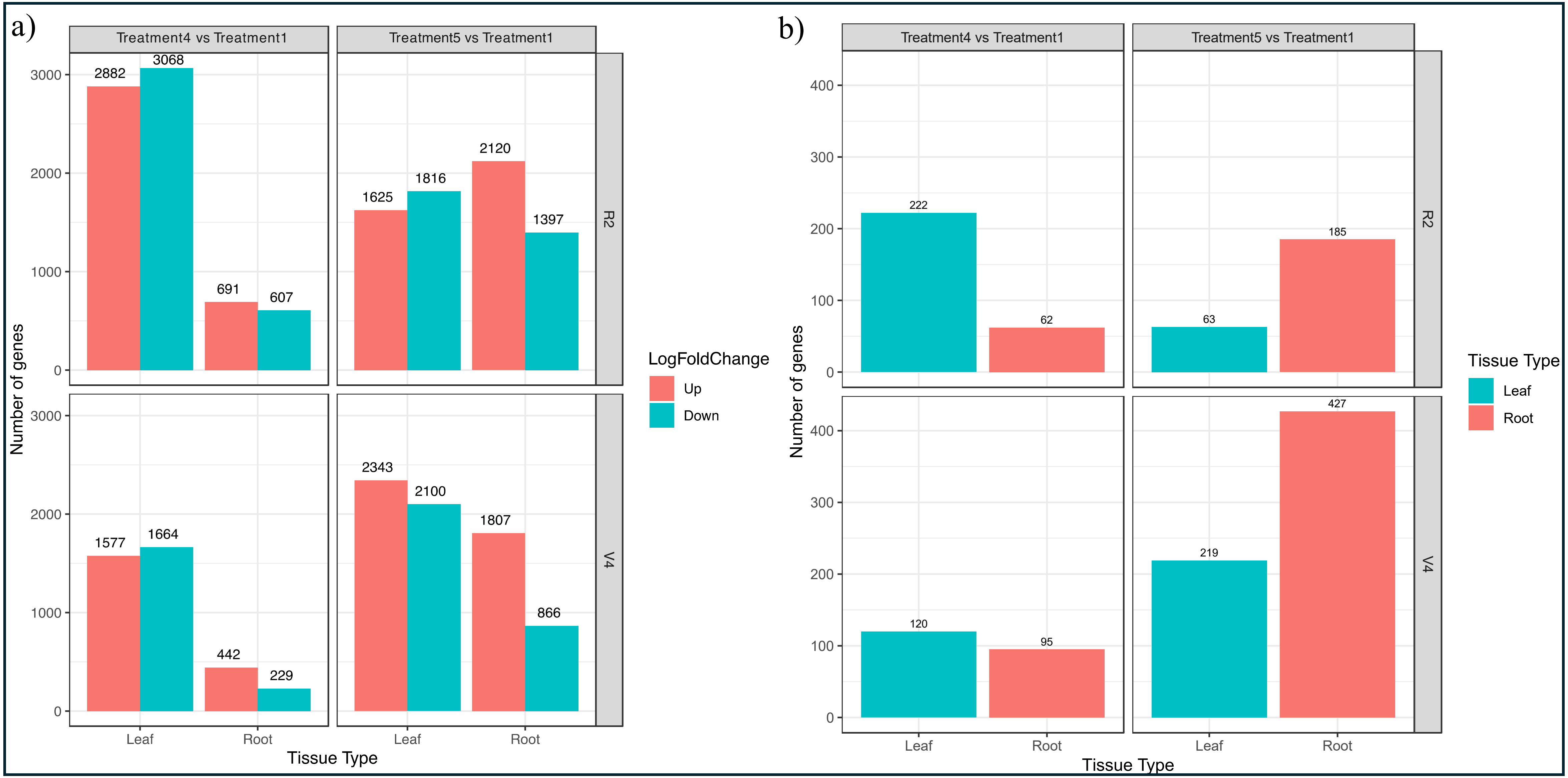


**Fig S2:** a) Differentially expressed genes (DEGs) are shown for each treatment and tissue type (Leaf and Root). The red bars represent upregulated genes, while the cyan bars represent downregulated genes. b) More upregulated DEGs (after applying $\log_2$fold change ≥ 2 threshold ) were observed in root tissue for both treatment 4 and 5 at the V4 and R2 stages. In contrast, more upregulated DEGs were found in leaf tissue at the V4 stage for treatment 5.

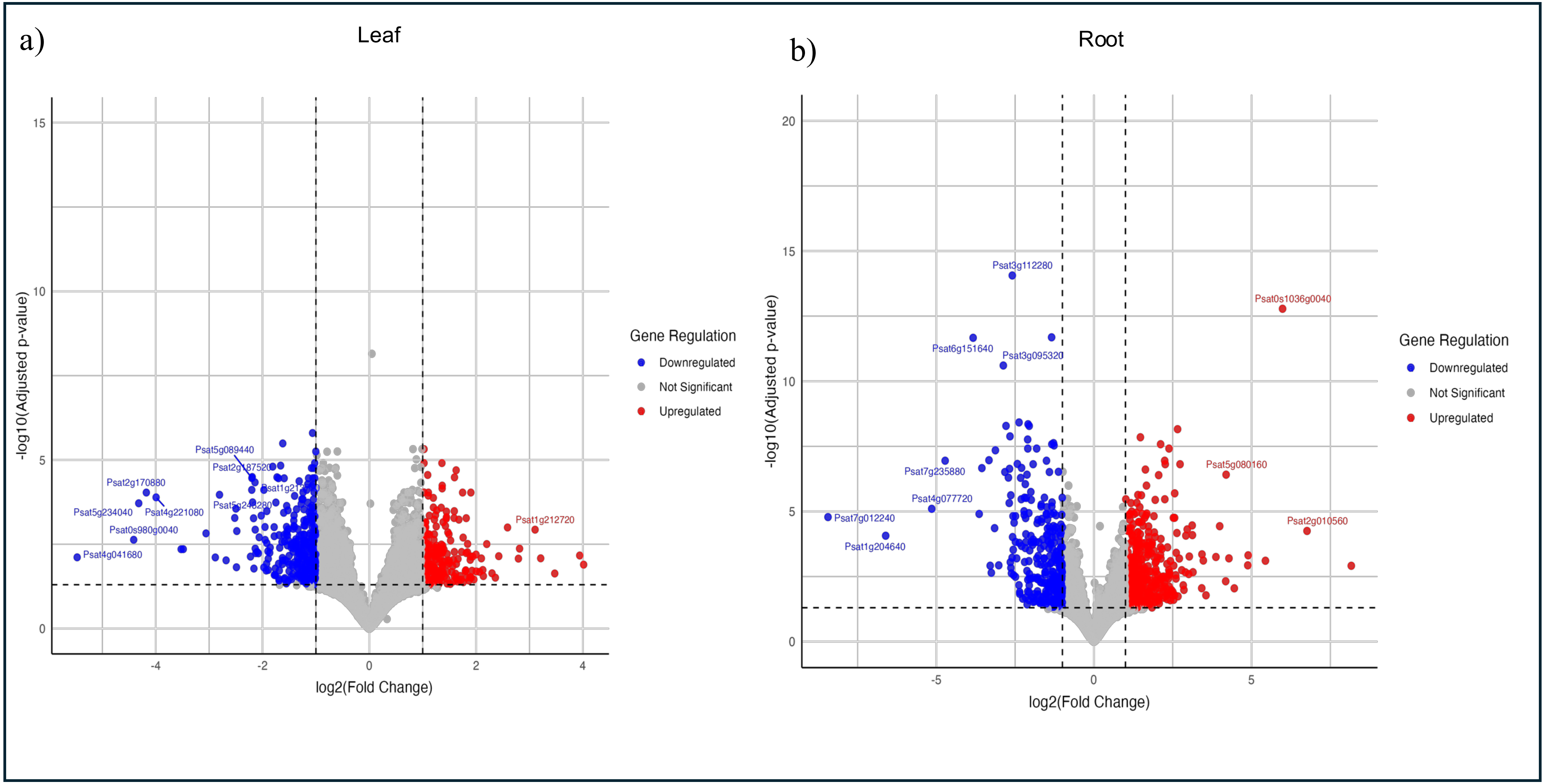


**Fig S3:** Volcano plot showing differentially expressed genes at the highest salinity treatment (Treatment 5) for a) Leaf and b) Root at reproductive (R2) stage. Genes with a $log_2$foldchange ≥ 2 threshold were considered upregulated.

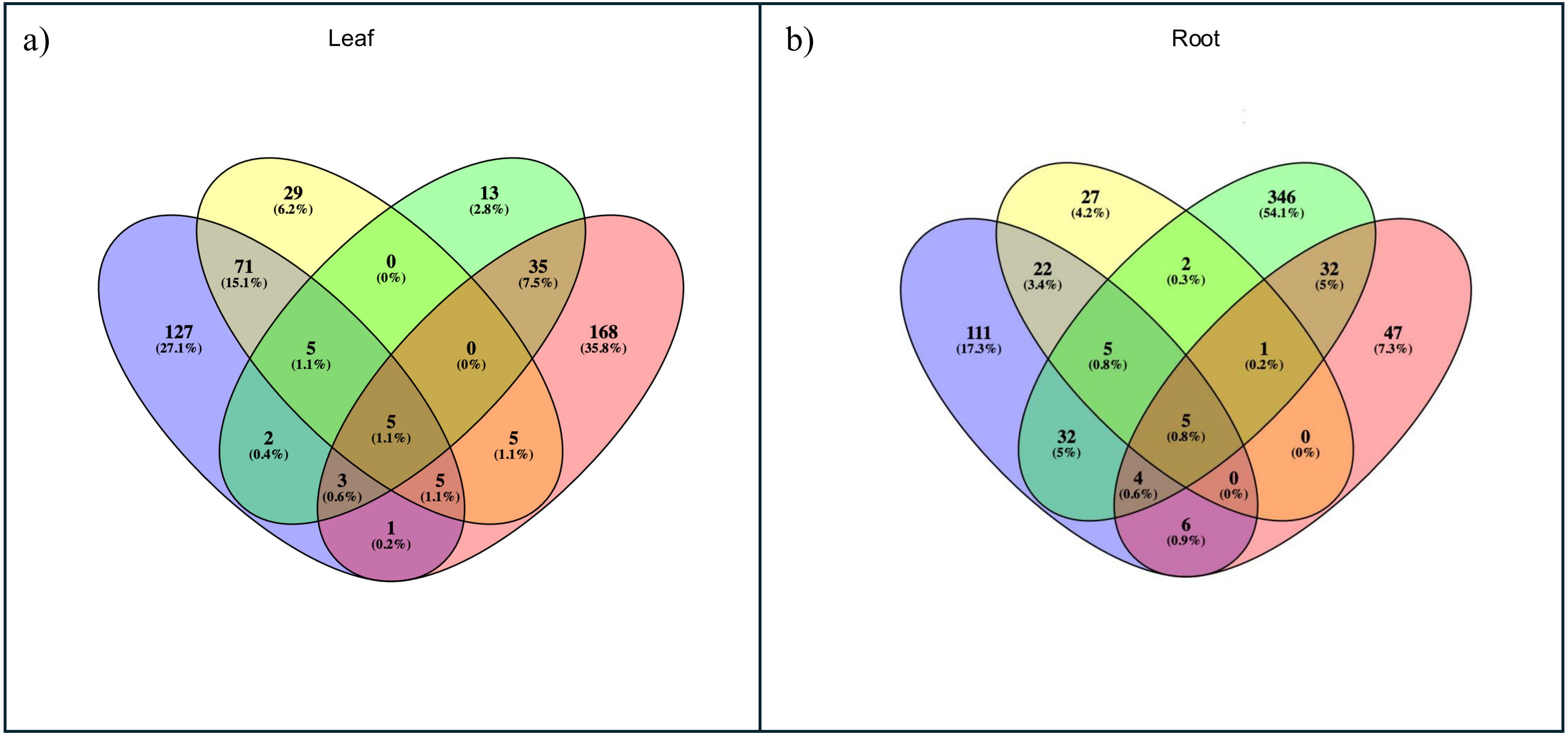


**Fig S4 :** Common DEGs among root and leaf tissue across all comparisons. a) Five genes were common across leaf tissues in treatment 4 and 5 at both vegetative (V4) and reproductive stages (R2). b) Five genes were common across root tissues in treatment 4 and 5 at both V4 and R2 stages.

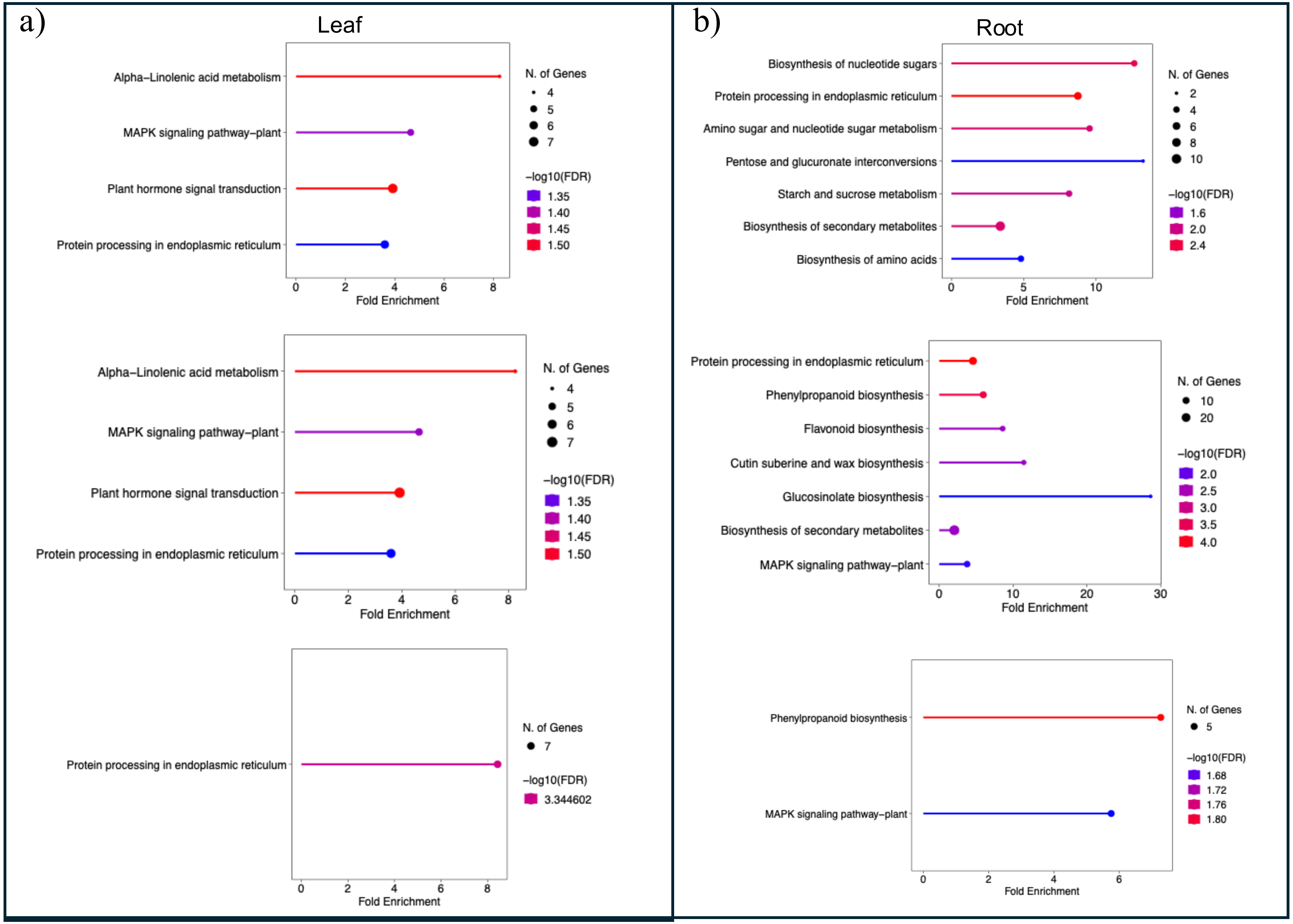


**Fig S5:** GO enrichment analysis for upregulated DEGs for a) Leaf and b) Root across treatments. Larger diameter circle in each plot represents total number of enriched genes.
